# The impact of genetic relationship between training and validation populations on genomic prediction accuracy in Atlantic salmon

**DOI:** 10.1101/2021.09.14.460263

**Authors:** Clémence Fraslin, José M. Yáñez, Diego Robledo, Ross D. Houston

## Abstract

The potential of genomic selection to improve production traits has been widely demonstrated in many aquaculture species. Atlantic salmon breeding programmes typically consist of sibling testing schemes, where traits that cannot be measured on the selection candidates are measured on the candidates’ siblings (such as disease resistance traits). While annual testing on close relatives is effective, it is expensive due to high genotyping and phenotyping costs. Therefore, accurate prediction of breeding values in distant relatives could significantly reduce the cost of genomic selection. The aims of this study were (i) to evaluate the impact of decreasing the genomic relationship between the training and validation populations on the accuracy of genomic prediction for two key target traits; body weight and resistance to sea lice; and (ii) to assess the interaction of genetic relationship with SNP density, a major determinant of genotyping costs. Phenotype and genotype data from two year classes of a commercial breeding population of Atlantic salmon were used. The accuracy of genomic predictions obtained within a year class was similar to that obtained combining the data from the two year classes for sea lice count (0.49 - 0.48) and body weight (0.63 - 0.61), but prediction accuracy was close to zero when the prediction was performed across year groups. Systematically reducing the relatedness between the training and validation populations within a year class resulted in decreasing accuracy of genomic prediction; when the training and validation populations were set up to contain no relatives with genomic relationships >0.3, the accuracies fell from 0.48 to 0.27 for sea lice count and from 0.63 to 0.29 for body weight. Lower relatedness between training and validation populations also tended to result in highly biased predictions. No clear interaction between decreasing SNP density and relatedness between training and validation population was found. These results confirm the importance of genetic relationships between training and selection candidate populations in salmon breeding programmes, and suggests that prediction across generations using existing approaches would severely compromise the efficacy of genomic selection.

## 1. Introduction

Genetic improvement of aquaculture species has a major and increasing role in providing sustainable seafood to meet the demands of a growing human population (Gjedrem, 2012). With increasing availability and affordability of genomic tools, molecular genetic markers can be routinely incorporated to improve the efficiency of aquaculture breeding programmes (Houston et al., 2020). The incorporation of such markers to improve prediction of breeding values for target traits occurs via two primary methods: marker-assisted selection and genomic selection. Marker-assisted selection has been successful for a limited number of traits where the genetic variation is controlled by major quantitative trait loci (QTL), e.g. resistance to Infectious Pancreatic Necrosis Virus (IPNV) in Atlantic salmon (Houston et al., 2008; Moen et al., 2009). Genomic selection (GS) is suitable for polygenic traits, and uses genome-wide genetic marker data to predict the genetic merit of the selection candidates (*i*.*e*. their breeding value) for target traits. In GS, genotype and phenotype data are typically collected in a training population and used to train a genomic prediction model, which is then used to predict the breeding values of selection candidates with genotype data only (Goddard and Hayes, 2007; Meuwissen et al., 2001). GS is routinely applied in advanced livestock and aquaculture breeding programmes, with notable benefits in terms of genetic gain and control of inbreeding (Boudry et al., 2021; Houston et al., 2020; You et al., 2020; Zenger et al., 2019). For any given trait, the accuracy of genomic prediction is highly dependent on the ability of the markers to accurately capture the genetic relationship between individuals from the training and selection candidate populations (Habier et al., 2007; Hayes et al., 2009; Villanueva et al., 2005). As such, in practice, the accuracy of genomic prediction is known to depend on the relationship between the training and selection populations (e.g. Wientjes et al., 2013).

With the recent development and availability of medium to high-density SNP arrays for most of the major aquaculture species (Griot et al., 2021; Houston et al., 2014; Liu et al., 2014; Palti et al., 2015; Peñaloza et al., 2021, 2020; Yáñez et al., 2014b), GS has begun to be widely applied in aquaculture breeding programmes. In recent years, both simulated and empirical data have shown that GS performs better than standard pedigree-based selection in test populations similar in structure to aquaculture breeding programs (Houston et al., 2020; Zenger et al., 2019). The high fecundity of aquaculture species enables the production of large full and half sibling families, which facilitates selection for traits that cannot be easily measured in the selection candidates, such as disease resistance or fillet quality, via their measurement on full- and half-siblings of the candidates.

The typical primary target of most aquaculture breeding programmes is growth rate, generally measured as the fish body weight or length. This trait can be easily measured throughout the life of the fish and has been reported to be moderate to highly heritable with a polygenic architecture (Baranski et al., 2010; Gutierrez et al., 2012; Sae-Lim et al., 2017; Tsai et al., 2015). Growth is easy to measure on the selection candidates themselves, however the rearing condition of the breeding nucleus can be quite different from the production environment resulting in different growth performance, thus, GS would be an efficient approach to select for improved growth in the production environment. Additionally, disease resistance traits are of the utmost importance in aquaculture breeding programs since disease outbreaks represent a major economic threat, and often few biosecurity and treatment options exist (Houston, 2017). Among the numerous pathogens threatening the Atlantic salmon industry, sea lice is probably the most important, a marine parasite causing millions of loses to the salmon industry worldwide (Abolofia et al., 2017; Costello, 2009), with *Caligus rogercresseyi* being the main species affecting the Southern Hemisphere, including Chile (Lhorente et al., 2019). Encouragingly, resistance to sea lice is moderately heritable and controlled by a polygenic architecture, and previous studies have shown the benefit of genomic selection over family selection (e.g. Correa et al., 2017a; Ødegård et al., 2014; Tsai et al., 2016).

However, genotyping large number of individuals using medium to high-density (HD) SNP platforms is still expensive, and therefore routine collection of genotype and phenotype data on large numbers of individuals each generation is expensive. In the past few years, a number of studies have focused on systematically testing low-density (LD) marker panels to help reduce the cost of GS (e.g. Lillehammer et al., 2013; Palaiokostas et al., 2019; Tsai et al., 2016; Tsairidou et al., 2020). Kriaridou et al., (2020) recently used four different datasets from four different species to demonstrate that SNP densities between 1,000 and 2,000 result in genomic prediction accuracies close to those obtained with HD panels. Previous studies in salmonid species suggest that between 1K and 20K SNPs are needed to reach genomic prediction accuracies close to those obtained using HD SNP panels in these species (Bangera et al., 2017; Correa et al., 2017a; Tsai et al., 2016; Yoshida et al., 2018a), with LD panels containing prioritised variants showing particular promise (Vallejo et al., 2018; Yoshida and Yáñez, 2021).

Most salmon breeding programmes rely on successive year classes composed of related individuals, and therefore combining information of two successive generations or “skipping” the data collection of one generation could be alternative strategies to reduce the cost of GS. While the impact of reducing the number of SNPs in GS prediction accuracy has been widely investigated, the impact of the genetic relationship between training and validation populations has not yet been widely studied in aquaculture species. Initial studies using Atlantic salmon (Tsai et al., 2016), rainbow trout (D’Ambrosio et al., 2020) and common carp (Palaiokostas et al., 2019) suggest that prediction accuracy drops dramatically as the relationship between training and validation populations becomes more distant. This is a scenario previously demonstrated in crop and livestock breeding (Clark et al., 2012; Habier et al., 2010). However, this has not yet been systematically studied in aquaculture species.

To assess the feasibility of potential new cost-effective GS strategies to improve traits of interest, the impact of different training population structures and genotyping strategies on the genomic prediction accuracy needs to be better understood. The aim of this study was to evaluate the impact of decreasing the genomic relationship between the training and validation populations on the accuracy of genomic prediction for two traits of major importance in Atlantic salmon breeding programs, and at varying SNP densities. Body weight and sea lice count data from two year classes of the same commercial breeding program were used to systematically test the effect of decreasing SNP density (from 33K to 100 SNPs) and decreasing genomic relatedness between fish from training and validation sets on the accuracy of genomic prediction.

## 2. Material and Methods

### 2.1. Fish production, infectious challenge and phenotyping

The Atlantic salmon (*Salmo salar*) population used in this study was composed of two year classes (2010 and 2014) from the breeding population of AquaChile (formerly Salmones Chaicas, X^th^ Region, Chile). The origin of this farmed Atlantic salmon population, as well as the establishment of the breeding program, including the introduction of ova to Chile for farming purposes, subsequent management and reproduction, breeding goal, and selection criteria are described in detail by Barria et al., (2018) and López et al., (2019). Details on reproduction tagging, rearing conditions, disease challenge and management of fish used in the present work are previously described in (Correa et al., 2017a, 2017b) for year class 2010 and (Robledo et al., 2019, 2018) for year class 2014. Fish from the two year classes are related as fish from year class 2010 are the aunts and uncles of fish from year class 2014.

Briefly, fish from both year classes were individually Passive Integrated transponder (PIT)-tagged, body length and weight were measured at different time points and fish were experimentally challenged with sea lice (*Caligus rogercresseyi*). Briefly, body weight (BW) and body length (BL) were recorded at tagging and at the end of the challenge for fish from year class 2010 and at the end and start of the disease challenge for fish from year class 2014. For each year class, fish were separated into three tanks and infestation with the parasite was carried out by depositing 13 to 24 (2010) or 50 (2014) lice per fish in the tank and stopping the water flow for 6h after infestation. Six (2010) or eight (2014) days after challenge, fish were euthanized, individually removed from the tank and the number of lice attached to the fins was counted under a magnifying lamp (recorded as sea lice count, SLC). A fin clip was taken from each fish for DNA extraction and genotyping.

### 2.2. Ethic statement

The challenge experiments and sampling procedures were performed under local and national regulatory systems and were approved by The Comité de Bioética Animal, Facultad de Ciencias Veterinarias y Pecuarias, Universidad de Chile (Santiago, Chile), under the certificate N° 08–2015 for fish from year class 2010 and the certificate N°01-2016 for fish from year class 2014. The Comité de Bioética Animal based its decision on the Council for International Organizations of Medical Sciences standards, in accordance with the Chilean standard NCh-324-2011.

### 2.3. Genotyping, imputation and quality controls

DNA from 2,404 and 2,668 fish for 2010 and 2014, respectively, was extracted from tissue samples using a commercial kit (Wizard R Genomic DNA Purification Kit, Promega) following manufacturer’s instructions. Fish from year class 2010 were genotyped using a custom 50K Affymetrix Axiom SNP array developed from a higher density (200K) SNP panel. The SNP discovery, filtering and construction of the 200K and 50K arrays are described in detail by Yáñez et al., (2016) and Correa et al., (2015), respectively. Fish from year class 2014 were genotyped using a custom-made 965 SNP panel and imputed to the same 50K SNPs of the 2010 year class using FImpute software (v2.2, Sargolzaei et al., 2014) as described in Robledo et al., (2019).

Standard quality control procedures were performed using the Plink software (version 1.9, Purcell et al., 2007) for the two year classes separately. Briefly, for year class 2010, SNPs with a call rate under 98%, a minor allele frequency (MAF) below 0.05 and deviating from Hardy-Weinberg equilibrium (p-value > 1.10^−6^) were removed from the dataset, resulting in 2,258 fish with an individual call rate over 95% and genotyped for 35,479 SNPs. For year class 2014, SNPs with a MAF below 0.05 and deviating from the Hardy-Weinberg equilibrium (p-value > 1.10^−6^) after imputation were removed from the dataset, resulting in 2,345 fish genotyped for 35,833 SNPs. Finally, only the 32,579 SNPs in common between the two year classes were retained in the dataset. Separately, the same quality controls were performed on the non-imputed low-density SNP panel of year class 2014, resulting in 2,345 fish with 873 SNPs.

### 2.4. Estimation of genetic parameters and genomic-based BLUP model

Variance components, heritability and genomic breeding values (GEBV) for both sea lice count (SLC) and body weight (BW) were estimated using the following linear mixed model:

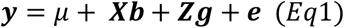

where ***y*** was the vector of phenotype (SLC or BW), *μ* is the overall mean of phenotypes, ***b*** is the vector of fixed effects and **X** the corresponding incidence matrix, ***g*** is the vector of random additive genetic effect following the normal distribution 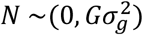 with 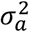 the additive genetic variance and **G** the genomic relationship matrix (GRM) as described in VanRaden (2008) and **Z** the corresponding incidence matrix. ***ε***_*i*_ is the vector of residual effects following the normal distribution 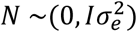with 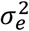 the residual variance and I the identity matrix. For year class 2010 the tank number was used as a fixed effect for both SLC and BW, and age at weighting (in days) was used as covariate for BW. For year class 2014, tank number was used as a fixed effect for both SLC and BW, initial body weight and age at recording (in days) were included as covariate for SLC and BW, respectively.

Genetic parameters were estimated by Average Information Restricted Maximum Likelihood algorithm (AI-REML) implemented in GCTA software (Yang et al., 2011). For this analysis, the GRM was built directly by GCTA with the following equation where the g_jk_ term of the matrix (genomic relationship between jth and kth fish) is estimated using the following equation:

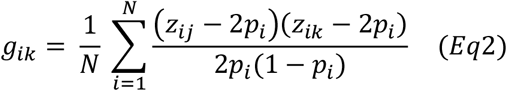

where N is the total number of SNP, z_ij_ and z_ik_ are numbers of copies of the reference allele for the i^th^ SNP for the j^th^ and k^th^ fish, respectively, and p_i_ is the frequency of the reference allele estimated from the markers.

GEBVs were estimated using the blupf90 program from BLUPf90 software (version 1.68, Misztal et al., 2002).

### 2.5. The Impact of Genetic Relationship on Genomic prediction

The accuracy of genomic prediction for resistance to sea lice as measure by sea lice counts (denoted SLC) and BW was only estimated for fish of year class 2014, and was assessed by replicates of a k-fold cross-validation (CV) procedure under four different scenarios (see below). For each scenario, the population was separated into k groups; one group was designated as the validation set and the phenotypes of the animals assigned to that group were masked, their genomic breeding values (GEBVs) were predicted from the remaining k-1 groups that composed the training set. The efficiency of genomic selection was assessed by the accuracy and bias of predicted GEBVs. The accuracy (r) of genomic prediction was calculated as the Pearson correlation coefficient between GEBVs and true phenotypes of the validation set fish divided by the square root of the trait heritability [*r* = *Corr*(*GEBV, y*)/*h*] (Legarra et al., 2008).

The selection bias (b) was estimated as the regression coefficient of the phenotypes on the predicted values. This coefficient is expected to be equal to 1 in the absence of bias. A coefficient below 1 indicates an over-dispersion of the GEBVs, on the contrary a coefficient below 1 indicates an under-dispersion of the GEBVs. The cross-validation process was replicated 10 or 20 times, depending on the scenario, and for each replicate a new randomization of the fish into k-groups was performed. For each scenario, the average and standard deviation of both the accuracy and bias were obtained for each k-fold and replicate.

To estimate the impact of the relationship between the training and the validation sets the following four scenarios were tested:

1. *Within year class 2014*. Five groups of equal size (n=469 fish per group) were created by randomly assigning fish from year class 2014 to groups using the CVrepGPAcalc package (v1.0 / R version 3.6.3) from (Tsairidou et al., 2020). The validation set was composed of fish from one group (n= 469, 20% of the population) with their phenotypic values masked and their GEBVs predicted using the genomic and phenotypic values of the training set comprising the remaining four groups (n=1,876, 80% of the population). This procedure was performed 10 times.
2. *Across year classes*. To assess the efficiency of using phenotypic values from a previous year class to predict the values of the next generation, the full dataset from year class 2010 was used as training set to predict the GEBVs of all fish from year class 2014.
3. *Combining both year classes*. To assess if combining cross-generation information improves prediction accuracy, fish from both year classes 2010 and 2014 were used to predict GEBVs of fish from year class 2014. The same groups as in scenario 1 were used. Within one replicate, one group from 2014 (n=469) was used as validation set and the remaining four groups from year class 2014 were merged with all the fish from year class 2010 and used as the training set (n=4,134 fish). This procedure was performed 10 times.
4. *Using genomic relationship threshold*. To assess the effect of the genomic relationship between training and validation sets within year class 2014, three groups of equal size were created so that the genomic relationship (obtained from the GRM estimated with GCTA software) between two fish assigned to two different groups was below a predefined kinship threshold. In this scenario, all fish with a genomic relationship above the predefined threshold could be assigned to the same group. Nine different genomic kinship thresholds were used: 0.3, 0.33, 0.35, 0.37, 0.4, 0.45, 0.5, 0.55 and no threshold. The GEBVs of fish from one group (1/3^rd^ of the population) were predicted using the genomic and phenotypic values of the remaining group (2/3^rd^ of the population). This procedure was performed 20 times.

The impact of reducing the SNP density on genomic prediction accuracy was tested in year class 2014 with nine randomly generated low to medium density SNP panels. For each panel, SNPs were randomly sampled from the 32K SNPs from the HD panel using the CVrepGPAcalc package (Tsairidou et al., 2020). The sampling was performed within each chromosome without replacement, with the number of SNP from a given chromosome being proportional to the physical length of the chromosome in the *S. salar* reference genome assembly (Lien et al., 2016, Genbank accession GCA_000233375.4). Because the number of SNPs was proportional to chromosome length, the total number of SNP selected to build a panel was allowed to differ slightly from the target density (Supplementary Table S1). For each target density, 10 panel replicates were generated, which were allowed to overlap by chance. Genetic parameters were estimated as described above (Eq1) with a new GRM built for each low-density panel.

The impact of reducing the SNP density was analysed within year class 2014 (scenario 1) and combined with the analysis of the genomic relationship between training and validation populations (scenario 4). For the latter, five different SNP density panels were tested (10K, 5K, 1K, 500, 100) for five kinship thresholds (0.3, 0.33, 0.35, 0.37 and 0.4).

### 2.6. Data availability

For fish from year class 2014, the imputed genotypes and corresponding SNP positions and phenotypes of the challenged animals are available, respectively, in the Supplementary Data Sheet 1 (compressed file, GenABEL.ped and GenABEL.map files) and in Supplementary Table 2 from Robledo et al., (2019). Phenotype and genotype data for fish from year class 2010 are available at https://figshare.com/articles/Compative_genomic_of_O_mykiss_and_S_salar_for_resistance_to_Sea_lice/7676147, from Cáceres et al., (2021).

## 3. Results and discussion

### 3.1. Genetic parameters estimates

Estimates of genetic parameters for the two traits and the two year classes are summarised in Table 1. Heritability estimates for SLC were low to moderate and quite different between the two year classes with a heritability of 0.11 (± 0.025 se) for year class 2010 and a heritability of 0.29 (± 0.035 se) for year class 2014. Those values were within the range of what has been previously reported for resistance to sea lice (0.10-0.27; (Cáceres et al., 2021; Correa et al., 2017a, 2017b; Ødegård et al., 2014; Tsai et al., 2016; Tsairidou et al., 2020; Yáñez et al., 2014a). For BW, heritability estimates were slightly higher, 0.31 (± 0.035 se) and 0.40 (± 0.035 se) for year classes 2010 and 2014, respectively, also in range with previous estimates. Previous studies reported pedigree based heritability estimates of 0.2-0.49 for BW (Gutierrez et al., 2015; Tsai et al., 2015; Yáñez et al., 2014a) and genomic based estimates of 0.27-0.6 for standardised or log transformed BW (Sae-Lim et al., 2017; Tsai et al., 2015; Tsairidou et al., 2020; Yoshida et al., 2017). It should be noted that the average number of lice per fish was substantially lower for fish from year class 2010 (5.12 ± 4.43 sd) than for fish from year class 2014 (39.0 ± 16.40 sd). As BW was measured at tagging for fish from year class 2010, average BW was significantly lower (13.3g ± 3.32 sd) for the older year class than for the fish from year class 2014 (122.1g ± 40.03 sd), measured prior to the sea lice challenge. These differences in the trait values between the year groups may affect the ability to predict breeding values across year groups, in addition to the impact of genetic relationship.

**Table 1.** Genetic parameters estimates for sea lice count and body weight for the two year class YC1 and YC2.

| Trait | Year class | Va (mean ± se) | Vp (mean ± se) | Ve (mean ± se) | h<sup>2</sup> (mean ± se) |
| --- | --- | --- | --- | --- | --- |
| SLC | 2010 | 1.41 ± 0.328 | 12.33 ± 0.389 | 10.92 ± 0.381 | 0.11 ± 0.025 |
| SLC | 2014 | 61.31 ± 8.478 | 213.36 ± 6.747 | 152.05 ± 7.238 | 0.29 ± 0.035 |
| BW | 2010 | 3.48 ± 0.490 | 11.17 ± 0.426 | 7.69 ± 0.312 | 0.31 ± 0.035 |
| BW | 2014 | 618.16 ± 66.518 | 1554.78 ± 51.063 | 936.62 ± 49.042 | 0.40 ± 0.035 |

SLC = sea lice count, BW = body weight measured at tagging for 2010 and prior to infection for 2014.
Vg = genetic variance, Ve = residual variance, Vp = phenotypic variance (Vg + Ve), h<sup>2</sup> = heritability estimated as Vg / (Vg + Ve).

### 3.2. Accuracy of genomic predictions within and across year class

The results of genomic selection for predictions (1) within year class 2014, (2) across year classes, and (3) combining both year classes using the full density SNP panel are summarised in Table 2. For both traits, the highest accuracy of genomic prediction was obtained when the training and validation sets were created with just animals of year class 2014 (1), but combining both year classes (3) resulted in practically the same accuracy (the decrease in accuracy was not significant, p-value = 0.55 for SLC and p-value = 0.18 for BW). These values were in the range of previously reported accuracies for similar SNP density panels in Atlantic salmon (Tsai et al., 2016, 2015) and slightly below those found by Yoshida et al., (2018b) for BW.

**Table 2.** Genomic prediction accuracy and bias for sea lice count and body weight in year class 2014 estimated under three scenarios using different training sets.

|  | 2014 predicted with 2014<br>(scenario 1) |  | 2014 predicted with<br>2010 (scenario 2) |  | 2014 predicted with 2010 and<br>2014 (scenario 3) |  |
| --- | --- | --- | --- | --- | --- | --- |
|  | Accuracy | Bias | Accuracy | Bias | Accuracy | Bias |
| **SLC** | 0.49 ± 0.087 | 0.99 ± 0.203 | 0.057 | 0.88 (0.593) | 0.48 ± 0.091 | 1.03 ± 0.222 |
| **BW** | 0.62 ± 0.067 | 1.03 ± 0.159 | - 0.033 | -1.16 (1.142) | 0.61 ± 0.068 | 0.83 ± 0.129 |

SLC = sea lice count, BW = body weight measured at tagging for 2010 and prior to infection for 2014.
Accuracy = Correlation (GEBV, phenotype) / h with SLC $h^2 = 0.287$ and BW $h^2 = 0.398$
Bias = regression coefficient of (phenotype ~ prediction)
Mean ± standard deviation of accuracy and bias over 5 folds and 10 cross validation sets.
For bias in scenario 2, standard error in brackets.

The accuracy of the genomic prediction from year class 2010 fish to 2014 fish (2) was close to zero. The low performance of scenario 2 and the fact that, despite almost doubling the training population size in scenario 3, no effect on the prediction accuracy for neither of the traits studied was observed, may reflect the relatively distant relationship between the two year classes, since the two generations are only second-degree relatives (year class 2010 was composed of uncles and aunts of fish from year class 2014). These results are consistent with the findings of Tsai et al., (2016), that estimated very low genomic prediction accuracies across two year groups of the same commercial population. Combining data of first-degree relatives across generations (e.g. direct parents of selection candidates) could result in a more efficient selection, but this would only be possible for traits that can be measured in the selection candidates, and therefore the benefit would be limited. The results may also reflect the aforementioned differences in the traits measured in the two year classes. Genetic correlations were estimated as low between the traits measured in the two year classes, albeit the structure of the data was not amenable to accurate assessment of this correlation.

Predictions obtained under scenarios 1 and 3 showed little evidence of bias whereas predictions obtained under scenario 2 resulted in highly biased prediction values for both traits (Table 2). Two previous publication by D’Ambrosio et al., (2020) for several female reproduction traits in rainbow trout and by Palaiokostas et al., (2019) for common carp resistance to Koi Herpesvirus disease, similarly reported that distantly related training and validation populations were also associated with highly biased predictions.

### 3.3. Impact of SNP density on prediction accuracy

Decreasing the SNP density used to build the GRM for the GBLUP analysis caused a decrease in the accuracy of genomic selection, with the lowest accuracy values (0.26 ± 0.096 for SLC, 0.27 ± 0.076) obtained for the lowest SNP density (100) (see Supplementary Figure S1). For both traits the genomic prediction accuracy was less than 10% lower when estimated with 3K SNPs compared to the full imputed 32K SNPs and it was about 6% lower for 5K SNPs compared with 32K. For SNP densities of 1K or lower the accuracy dropped and was at least 20% lower. These results are in agreement with previous studies in salmonid species, which indicate that a range between 1K to 20K are needed to reach accuracies of genomic predictions close to those obtained using HD SNP panels (Bangera et al., 2017; Correa et al., 2017a; Tsai et al., 2016; Vallejo et al., 2018; Yoshida et al., 2018a). SNP panels with densities lower than 3K can also be applied without losing any accuracy when prioritizing variants based on their effect on a particular trait (Vallejo et al., 2018; Yoshida and Yáñez, 2021). As previously reported in Tsairidou et al., (2020) the variability in prediction accuracy between SNP panel replicates was substantially larger at lower SNP densities. Variation patterns were similar between the two traits with the exception of the accuracy of the 1K density panels, which was more variable for SLC than for BW (Supplementary Figure S1). However, regardless of the SNP density of the panel used, the genomic predictions did not show any sign of bias (Supplementary Table S2).

### 3.4. Impact of genomic relationship on prediction accuracy

The relationship between training and validation populations appears to be critical for efficient genomic selection but, to the best of our knowledge, its impact has never been systematically tested within a typical aquaculture population. The impact of progressively decreasing the relationship between training and validation sets on the accuracy and bias of genomic prediction (scenario 4) for both SLC and BW are presented in Figure 1. In this study, genomic relationship thresholds were used to systematically exclude close relationships from the training and validation populations. Genomic relationship thresholds from 0.55 to 0.3 were tested. Due to the family structure of the population, it was not possible to reduce the threshold between fish in the training and validation sets below 0.3. As expected, when the training and validation sets were less related the genomic predictions were less accurate for both traits. When the genomic relationship threshold was set at 0.4 or higher (*i*.*e*. equivalent to a full-sib relationship) the accuracy of genomic prediction was similar to the accuracy that can be expected for the trait based on a random cross validation set (no kinship threshold). With genomic relationship threshold between the two sets equal or below 0.37, the prediction accuracy started to decrease dramatically, reaching a minimum when the genomic relationship between training and validation sets was 0.3 (approximately equivalent to the relationship between half-siblings). When the relationship threshold between training and validation sets was 0.37, the accuracy of genomic prediction was only reduced by 4% for BW whereas it was reduced by 12% for SLC. When the relationship threshold was 0.3, the accuracy of genomic prediction was 44% lower for SLC (0.27 ± 0.048) and 51% lower for BW (0.30 ± 0.107).

**Figure 1.**
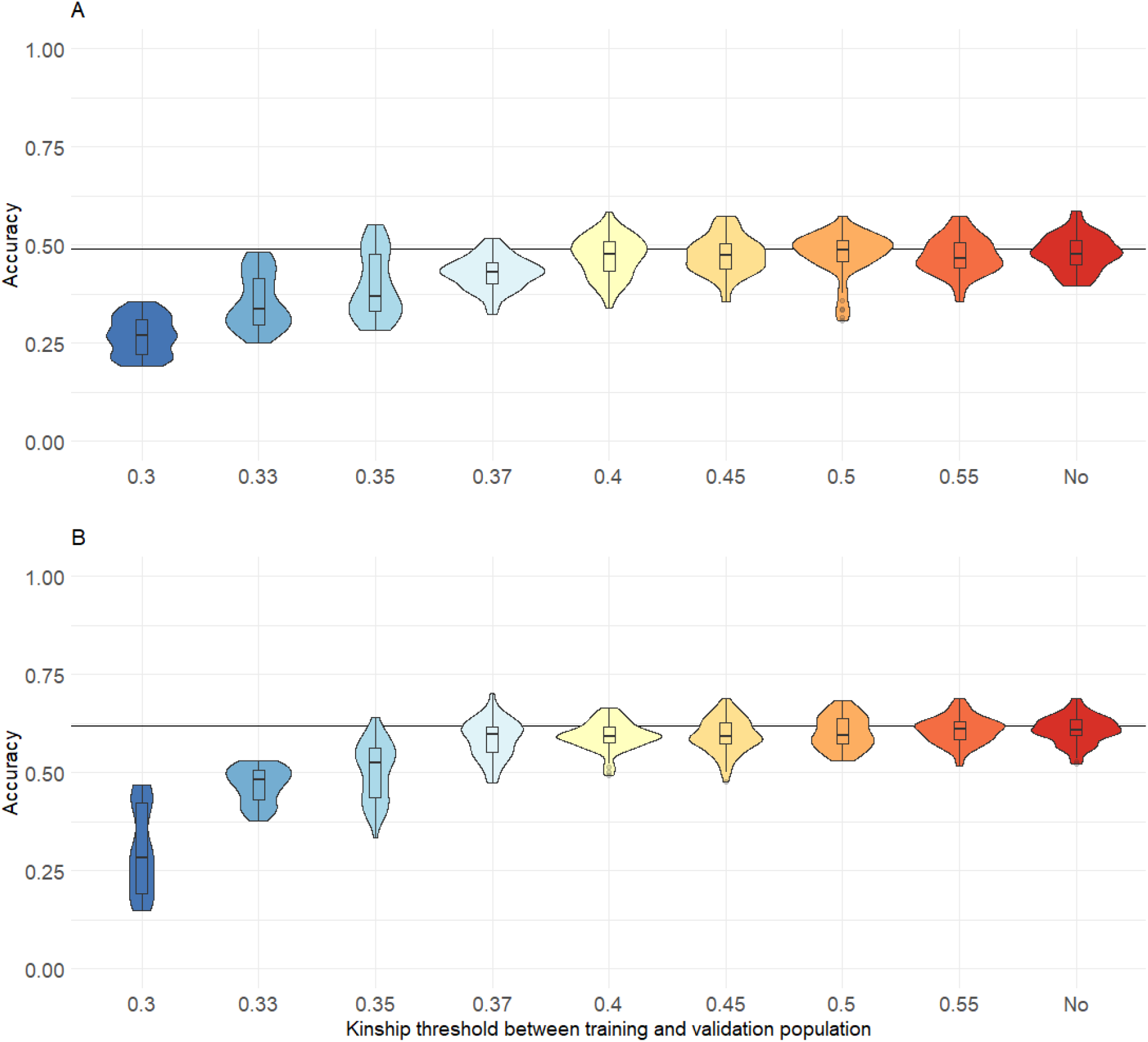
Accuracy of genomic prediction for sea lice count (A) and body weight (B) within the generation 2014 estimated with decreasing values of genomic relationship between training and validation sets. The dark line represent the mean accuracy of GBLUP obtained with random cross validation sets (10 simulation 5 groups). Fish from year class 2014 were assigned to three groups according to their genomic relationship in order to keep the genomic relationship between individuals of two different groups below a certain kinship threshold.

While decreasing the SNP density did not seem to result in increased prediction bias, decreasing the degree of relationship between training and validation populations did induce an over-dispersion of the variance of the GEBVs, with a bias of 0.73 (± 0.231) for SLC and 0.69 (± 0.232) for BW (See Supplementary Table S2) for the lowest genomic kinship threshold. Interestingly, for BW the predictions were less biased (value closer to 1) with a kinship degree threshold of 0.33, whereas for that same threshold SLC prediction variance was still overestimated (bias of 0.84). Those results are in accordance with a genomic selection study in common carp where Palaiokostas et al., (2019) tested several scenarios based on pedigree relationship. When only half-sibs of the selection candidates were included in the training set, they observed a small decrease (6-8%) of the genomic prediction accuracy, but predictions based on non-sibs (*i*.*e*. separate families) highly reduced the accuracy (up to a 72% decrease).

### 3.5. Interaction between SNP density and genetic relationship

The interaction between reduced density SNP panels and the genetic relationship between training and validation sets was also investigated (Figures 2A and 2B, supplementary table S4). In this scenario, the accuracy obtained with the highest density panel (10K SNPs) and lowest (0.3) genomic relationship was in the same range (0.25 ± 0.050 for SLC, 0.30 ± 0.103 for BW) as the accuracy obtained with the smallest density panel (100 SNPs) and highest (0.4) genomic relationship (0.25 ± 0.068 for SLC, 0.25 ± 0.060 for BW). For SLC, regardless of the tested SNP density, when the genomic relationship threshold between the training and validation sets was reduced from 0.4 to 0.3, the accuracy decreased by 49.4% on average. Whereas, when comparing accuracy between the highest and the lowest SNP density (10K vs 100 SNPs), accuracy decreased by 47.5% on average across all genomic similarity thresholds. For body weight, the decrease in accuracy was more striking when the SNP density decreased (64.9% on average across genomic relationship for 10K vs 100 SNPs), than when the genomic relationship decreased (58.8% on average for 0.4 vs 0.3 genomic relationship). The simultaneous decrease of both parameters resulted in a major drop in accuracy, which was reduced by 74% for SLC and by 89% for BW with 100 SNPs and a genomic relationship threshold between training and validation sets of 0.3.

**Figure 2.**
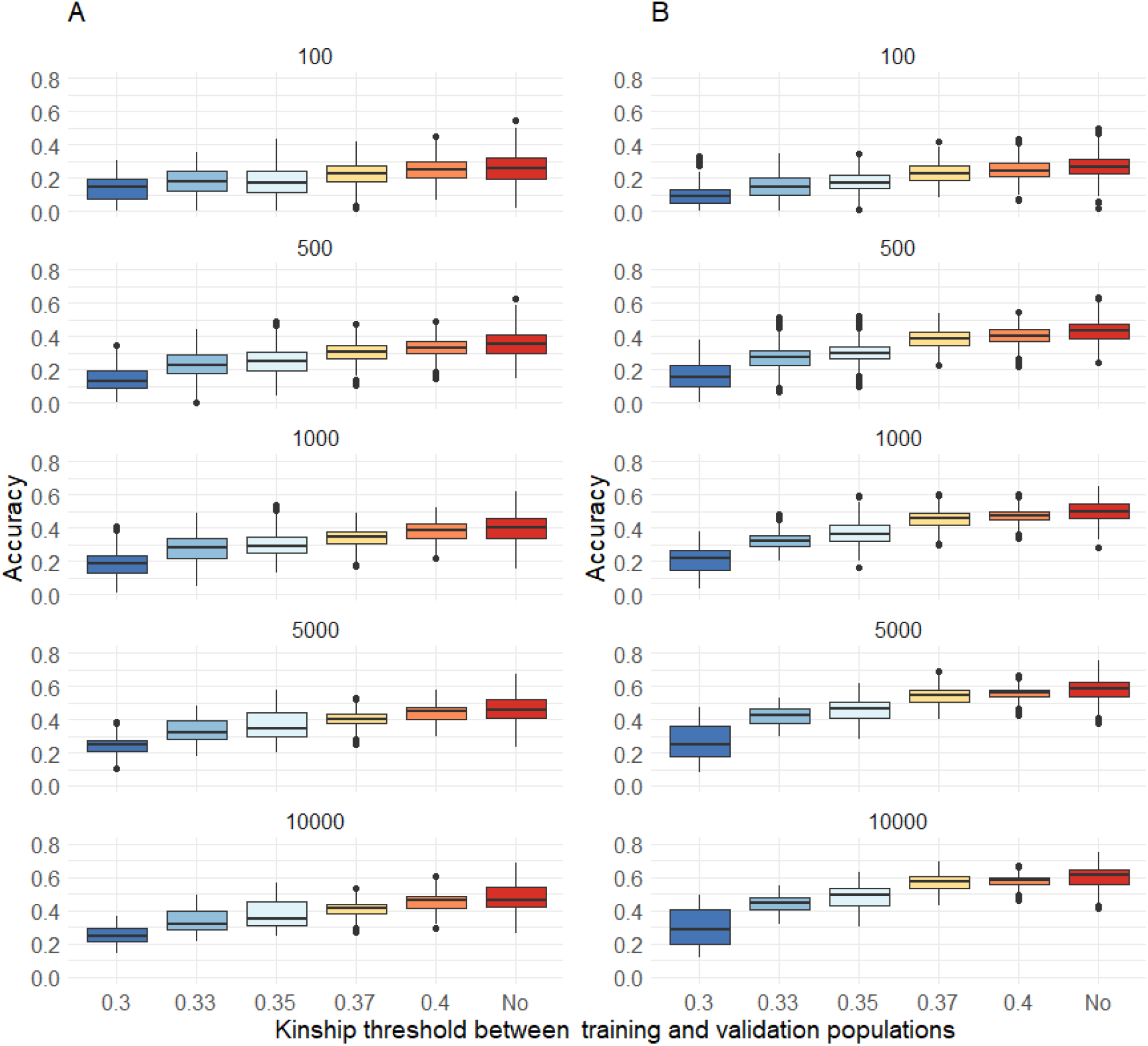
Accuracy of genomic prediction for sea lice count (A) and initial body weight (B) estimated with different SNP density panels and genomic relationship between training and validation sets. Box plot of accuracy of genomic prediction (GBLUP), estimated using various SNP density panels after 20 simulations with three cross-validation groups constructed to keep the genomic relationship between individuals of two different groups below a threshold, for sea lice count (A) or initial body weight (B).

When the relationship degree between training and validation populations was the lowest (0.3 or 0.33), predictions for both traits were highly biased regardless of the SNP density (Supplementary table S5), with the most extreme bias values (variance of GEBV highly overestimated) obtained for the lowest densities.

## 4. Conclusion

In this study, the impact of systematically decreasing the genomic relationship between training and validation populations on the accuracy of genomic prediction was tested for two traits of major importance in Atlantic salmon breeding programs. The interaction between genomic relationship threshold and varying SNP densities was also tested. Decreasing the relationship between the training and the validation population resulted in less accurate and more biased genomic prediction, which confirms the importance of building a testing population that contains close relatives (*i*.*e*. full and half siblings) of the selection candidates. Although there was no clear interaction between decreasing SNP density and relatedness between training and validation population, the simultaneous decrease of both parameters resulted in a major drop in accuracy. Therefore, the use of low density markers panels for cost-effective selection, although appropriate when genomic relationships are high, should be considered with care when genomic relationships are more distant.

## Acknowledgment

We would like to thanks the contribution of Benchmark Genetics Chile and AquaChile for providing the biological material and phenotypic records of the experimental challenge.

## Author contribution

RDH, DR, and JMY were responsible for the concept and design of this work. CF performed bioinformatics and statistical analyses. CF, RH, DR, and JMY drafted the manuscript. All authors read and approved the final manuscript.

## References

Abolofia, J., Asche, F., Wilen, J.E., 2017. The Cost of Lice: Quantifying the Impacts of Parasitic Sea Lice on Farmed Salmon. Mar. Resour. Econ. 32, 329–349. https://doi.org/10.1086/691981

Bangera, R., Correa, K., Lhorente, J.P., Figueroa, R., Yáñez, J.M., 2017. Genomic predictions can accelerate selection for resistance against Piscirickettsia salmonis in Atlantic salmon (Salmo salar). BMC Genomics 18, 121. https://doi.org/10.1186/s12864-017-3487-y

Baranski, M., Moen, T., Våge, D.I., 2010. Mapping of quantitative trait loci for flesh colour and growth traits in Atlantic salmon (Salmo salar). Genet. Sel. Evol. GSE 42, 17. https://doi.org/10.1186/1297-9686-42-17

Barria, A., López, M.E., Yoshida, G., Carvalheiro, R., Lhorente, J.P., Yáñez, J.M., 2018. Population Genomic Structure and Genome-Wide Linkage Disequilibrium in Farmed Atlantic Salmon (Salmo salar L.) Using Dense SNP Genotypes. Front. Genet. 9, 649. https://doi.org/10.3389/fgene.2018.00649

Boudry, P., Allal, F., Aslam, M.L., Bargelloni, L., Bean, T.P., Brard-Fudulea, S., Brieuc, M.S.O., Calboli, F.C.F., Gilbey, J., Haffray, P., Lamy, J.-B., Morvezen, R., Purcell, C., Prodöhl, P.A., Vandeputte, M., Waldbieser, G.C., Sonesson, A.K., Houston, R.D., 2021. Current status and potential of genomic selection to improve selective breeding in the main aquaculture species of International Council for the Exploration of the Sea (ICES) member countries. Aquac. Rep. 20, 100700. https://doi.org/10.1016/j.aqrep.2021.100700

Cáceres, P., Barría, A., Christensen, K.A., Bassini, L.N., Correa, K., Garcia, B., Lhorente, J.P., Yáñez, J.M., 2021. Genome-scale comparative analysis for host resistance against sea lice between Atlantic salmon and rainbow trout. Sci. Rep. 11, 13231. https://doi.org/10.1038/s41598-021-92425-3

Clark, S.A., Hickey, J.M., Daetwyler, H.D., van der Werf, J.H., 2012. The importance of information on relatives for the prediction of genomic breeding values and the implications for the makeup of reference data sets in livestock breeding schemes. Genet. Sel. Evol. 44, 4. https://doi.org/10.1186/1297-9686-44-4

Correa, K., Bangera, R., Figueroa, R., Lhorente, J.P., Yáñez, J.M., 2017a. The use of genomic information increases the accuracy of breeding value predictions for sea louse (Caligus rogercresseyi) resistance in Atlantic salmon (Salmo salar). Genet. Sel. Evol. 49, 15. https://doi.org/10.1186/s12711-017-0291-8

Correa, K., Lhorente, J.P., Bassini, L., López, M.E., Di Genova, A., Maass, A., Davidson, W.S., Yáñez, J.M., 2017b. Genome wide association study for resistance to Caligus rogercresseyi in Atlantic salmon (Salmo salar L.) using a 50K SNP genotyping array. Aquaculture, International Symposium on Genetics in Aquaculture XII (ISGA XII) 472, 61–65. https://doi.org/10.1016/j.aquaculture.2016.04.008

Correa, K., Lhorente, J.P., López, M.E., Bassini, L., Naswa, S., Deeb, N., Di Genova, A., Maass, A., Davidson, W.S., Yáñez, J.M., 2015. Genome-wide association analysis reveals loci associated with resistance against Piscirickettsia salmonis in two Atlantic salmon (Salmo salar L.) chromosomes. BMC Genomics 16, 854. https://doi.org/10.1186/s12864-015-2038-7

Costello, M.J., 2009. The global economic cost of sea lice to the salmonid farming industry. J. Fish Dis. 32, 115–118. https://doi.org/10.1111/j.1365-2761.2008.01011.x

D’Ambrosio, J., Morvezen, R., Brard-Fudulea, S., Bestin, A., Acin Perez, A., Guéméné, D., Poncet, C., Haffray, P., Dupont-Nivet, M., Phocas, F., 2020. Genetic architecture and genomic selection of female reproduction traits in rainbow trout. BMC Genomics 21, 558. https://doi.org/10.1186/s12864-020-06955-7

Gjedrem, T., 2012. Genetic improvement for the development of efficient global aquaculture: A personal opinion review. Aquaculture Complete, 12–22. https://doi.org/10.1016/j.aquaculture.2012.03.003

Goddard, M. e., Hayes, B. j., 2007. Genomic selection. J. Anim. Breed. Genet. 124, 323–330. https://doi.org/10.1111/j.1439-0388.2007.00702.x

Griot, R., Allal, F., Phocas, F., Brard-Fudulea, S., Morvezen, R., Bestin, A., Haffray, P., François, Y., Morin, T., Poncet, C., Vergnet, A., Cariou, S., Brunier, J., Bruant, J.-S., Peyrou, B., Gagnaire, P.-A., Vandeputte, M., 2021. Genome-wide association studies for resistance to viral nervous necrosis in three populations of European sea bass (Dicentrarchus labrax) using a novel 57k SNP array DlabChip. Aquaculture 530, 735930. https://doi.org/10.1016/j.aquaculture.2020.735930

Gutierrez, A.P., Lubieniecki, K.P., Davidson, E.A., Lien, S., Kent, M.P., Fukui, S., Withler, R.E., Swift, B., Davidson, W.S., 2012. Genetic mapping of quantitative trait loci (QTL) for body-weight in Atlantic salmon (Salmo salar) using a 6.5K SNP array. Aquaculture 358–359, 61–70. https://doi.org/10.1016/j.aquaculture.2012.06.017

Gutierrez, A.P., Yáñez, J.M., Fukui, S., Swift, B., Davidson, W.S., 2015. Genome-wide association study (GWAS) for growth rate and age at sexual maturation in Atlantic salmon (Salmo salar). PloS One 10, e0119730. https://doi.org/10.1371/journal.pone.0119730

Habier, D., Fernando, R.L., Dekkers, J.C.M., 2007. The Impact of Genetic Relationship Information on Genome-Assisted Breeding Values. Genetics 177, 2389–2397. https://doi.org/10.1534/genetics.107.081190

Habier, D., Tetens, J., Seefried, F.-R., Lichtner, P., Thaller, G., 2010. The impact of genetic relationship information on genomic breeding values in German Holstein cattle. Genet. Sel. Evol. 42, 5. https://doi.org/10.1186/1297-9686-42-5

Hayes, B.J., Visscher, P.M., Goddard, M.E., 2009. Increased accuracy of artificial selection by using the realized relationship matrix. Genet. Res. 91, 47–60. https://doi.org/10.1017/S0016672308009981

Houston, R.D., 2017. Future directions in breeding for disease resistance in aquaculture species. Rev. Bras. Zootec. 46, 545–551. https://doi.org/10.1590/s1806-92902017000600010

Houston, R.D., Bean, T.P., Macqueen, D.J., Gundappa, M.K., Jin, Y.H., Jenkins, T.L., Selly, S.L.C., Martin, S.A.M., Stevens, J.R., Santos, E.M., Davie, A., Robledo, D., 2020. Harnessing genomics to fast-track genetic improvement in aquaculture. Nat. Rev. Genet. 1–21. https://doi.org/10.1038/s41576-020-0227-y

Houston, R.D., Haley, C.S., Hamilton, A., Guy, D.R., Tinch, A.E., Taggart, J.B., McAndrew, B.J., Bishop, S.C., 2008. Major quantitative trait loci affect resistance to infectious pancreatic necrosis in Atlantic salmon (Salmo salar). Genetics 178, 1109–1115. https://doi.org/10.1534/genetics.107.082974

Houston, R.D., Taggart, J.B., Cézard, T., Bekaert, M., Lowe, N.R., Downing, A., Talbot, R., Bishop, S.C., Archibald, A.L., Bron, J.E., Penman, D.J., Davassi, A., Brew, F., Tinch, A.E., Gharbi, K., Hamilton, A., 2014. Development and validation of a high density SNP genotyping array for Atlantic salmon (Salmo salar). BMC Genomics 15, 90. https://doi.org/10.1186/1471-2164-15-90

Kriaridou, C., Tsairidou, S., Houston, R.D., Robledo, D., 2020. Genomic Prediction Using Low Density Marker Panels in Aquaculture: Performance Across Species, Traits, and Genotyping Platforms. Front. Genet. 11. https://doi.org/10.3389/fgene.2020.00124

Legarra, A., Robert-Granié, C., Manfredi, E., Elsen, J.-M., 2008. Performance of Genomic Selection in Mice. Genetics 180, 611–618. https://doi.org/10.1534/genetics.108.088575

Lhorente, J.P., Araneda, M., Neira, R., Yáñez, J.M., 2019. Advances in genetic improvement for salmon and trout aquaculture: the Chilean situation and prospects. Rev. Aquac. 11. https://doi.org/10.1111/raq.12335

Lien, S., Koop, B.F., Sandve, S.R., Miller, J.R., Kent, M.P., Nome, T., Hvidsten, T.R., Leong, J.S., Minkley, D.R., Zimin, A., Grammes, F., Grove, H., Gjuvsland, A., Walenz, B., Hermansen, R.A., von Schalburg, K., Rondeau, E.B., Di Genova, A., Samy, J.K.A., Olav Vik, J., Vigeland, M.D., Caler, L., Grimholt, U., Jentoft, S., Inge Våge, D., de Jong, P., Moen, T., Baranski, M., Palti, Y., Smith, D.R., Yorke, J.A., Nederbragt, A.J., Tooming-Klunderud, A., Jakobsen, K.S., Jiang, X., Fan, D., Hu, Y., Liberles, D.A., Vidal, R., Iturra, P., Jones, S.J.M., Jonassen, I., Maass, A., Omholt, S.W., Davidson, W.S., 2016. The Atlantic salmon genome provides insights into rediploidization. Nature 533, 200–205. https://doi.org/10.1038/nature17164

Lillehammer, M., Meuwissen, T.H.E., Sonesson, A.K., 2013. A low-marker density implementation of genomic selection in aquaculture using within-family genomic breeding values. Genet. Sel. Evol. 45, 39. https://doi.org/10.1186/1297-9686-45-39

Liu, S., Sun, L., Li, Y., Sun, F., Jiang, Y., Zhang, Y., Zhang, J., Feng, J., Kaltenboeck, L., Kucuktas, H., Liu, Z., 2014. Development of the catfish 250K SNP array for genome-wide association studies. BMC Res. Notes 7, 135. https://doi.org/10.1186/1756-0500-7-135

López, M.E., Linderoth, T., Norris, A., Lhorente, J.P., Neira, R., Yáñez, J.M., 2019. Multiple Selection Signatures in Farmed Atlantic Salmon Adapted to Different Environments Across Hemispheres. Front. Genet. 10, 901. https://doi.org/10.3389/fgene.2019.00901

Meuwissen, T.H., Hayes, B.J., Goddard, M.E., 2001. Prediction of total genetic value using genome-wide dense marker maps. Genetics 157, 1819–1829.

Misztal, I., Tsuruta, S., Strabel, T., Auvray, B., Druet, T., Lee, D.H., 2002. BLUPF90 AND RELATED PROGRAMS (BGF90). Presented at the 7th World Congress on Genetics Applied to Livestock Production, Montpellier, France, p. 2.

Moen, T., Baranski, M., Sonesson, A.K., Kjøglum, S., 2009. Confirmation and fine-mapping of a major QTL for resistance to infectious pancreatic necrosis in Atlantic salmon (Salmo salar): population-level associations between markers and trait. BMC Genomics 10, 368. https://doi.org/10.1186/1471-2164-10-368

Ødegård, J., Moen, T., Santi, N., Korsvoll, S.A., Kjøglum, S., Meuwissen, T.H.E., 2014. Genomic prediction in an admixed population of Atlantic salmon (Salmo salar). Front. Genet. 5. https://doi.org/10.3389/fgene.2014.00402

Palaiokostas, C., Vesely, T., Kocour, M., Prchal, M., Pokorova, D., Piackova, V., Pojezdal, L., Houston, R.D., 2019. Optimizing Genomic Prediction of Host Resistance to Koi Herpesvirus Disease in Carp. Front. Genet. 10. https://doi.org/10.3389/fgene.2019.00543

Palti, Y., Gao, G., Liu, S., Kent, M.P., Lien, S., Miller, M.R., Rexroad, C.E., Moen, T., 2015. The development and characterization of a 57K single nucleotide polymorphism array for rainbow trout. Mol. Ecol. Resour. 15, 662–672. https://doi.org/10.1111/1755-0998.12337

Peñaloza, C., Manousaki, T., Franch, R., Tsakogiannis, A., Sonesson, A.K., Aslam, M.L., Allal, F., Bargelloni, L., Houston, R.D., Tsigenopoulos, C.S., 2021. Development and testing of a combined species SNP array for the European seabass (Dicentrarchus labrax) and gilthead seabream (Sparus aurata). Genomics 113, 2096–2107. https://doi.org/10.1016/j.ygeno.2021.04.038

Peñaloza, C., Robledo, D., Barría, A., Tr?nh, T.Q., Mahmuddin, M., Wiener, P., Benzie, J.A.H., Houston, R.D., 2020. Development and Validation of an Open Access SNP Array for Nile Tilapia (Oreochromis niloticus). G3 Genes Genomes Genet. 10, 2777–2785. https://doi.org/10.1534/g3.120.401343

Purcell, S., Neale, B., Todd-Brown, K., Thomas, L., Ferreira, M.A.R., Bender, D., Maller, J., Sklar, P., de Bakker, P.I.W., Daly, M.J., Sham, P.C., 2007. PLINK: A Tool Set for Whole-Genome Association and Population-Based Linkage Analyses. Am. J. Hum. Genet. 81, 559–575. https://doi.org/10.1086/519795

Robledo, D., Gutiérrez, A.P., Barría, A., Lhorente, J.P., Houston, R.D., Yáñez, J.M., 2019. Discovery and Functional Annotation of Quantitative Trait Loci Affecting Resistance to Sea Lice in Atlantic Salmon. Front. Genet. 10, 56. https://doi.org/10.3389/fgene.2019.00056

Robledo, D., Gutiérrez, A.P., Barría, A., Yáñez, J.M., Houston, R.D., 2018. Gene Expression Response to Sea Lice in Atlantic Salmon Skin: RNA Sequencing Comparison Between Resistant and Susceptible Animals. Front. Genet. 9, 287. https://doi.org/10.3389/fgene.2018.00287

Sae-Lim, P., Kause, A., Lillehammer, M., Mulder, H.A., 2017. Estimation of breeding values for uniformity of growth in Atlantic salmon (Salmo salar) using pedigree relationships or single-step genomic evaluation. Genet. Sel. Evol. 49, 33. https://doi.org/10.1186/s12711-017-0308-3

Sargolzaei, M., Chesnais, J.P., Schenkel, F.S., 2014. A new approach for efficient genotype imputation using information from relatives. BMC Genomics 15. https://doi.org/10.1186/1471-2164-15-478

Tsai, H.-Y., Hamilton, A., Tinch, A.E., Guy, D.R., Bron, J.E., Taggart, J.B., Gharbi, K., Stear, M., Matika, O., Pong-Wong, R., Bishop, S.C., Houston, R.D., 2016. Genomic prediction of host resistance to sea lice in farmed Atlantic salmon populations. Genet. Sel. Evol. GSE 48, 47. https://doi.org/10.1186/s12711-016-0226-9

Tsai, H.-Y., Hamilton, A., Tinch, A.E., Guy, D.R., Gharbi, K., Stear, M.J., Matika, O., Bishop, S.C., Houston, R.D., 2015. Genome wide association and genomic prediction for growth traits in juvenile farmed Atlantic salmon using a high density SNP array. BMC Genomics 16, 969. https://doi.org/10.1186/s12864-015-2117-9

Tsairidou, S., Hamilton, A., Robledo, D., Bron, J.E., Houston, R.D., 2020. Optimizing Low-Cost Genotyping and Imputation Strategies for Genomic Selection in Atlantic Salmon. G3 GenesGenomesGenetics 10, 581–590. https://doi.org/10.1534/g3.119.400800

Vallejo, R.L., Silva, R.M.O., Evenhuis, J.P., Gao, G., Liu, S., Parsons, J.E., Martin, K.E., Wiens, G.D., Lourenco, D.A.L., Leeds, T.D., Palti, Y., 2018. Accurate genomic predictions for BCWD resistance in rainbow trout are achieved using low-density SNP panels: Evidence that long-range LD is a major contributing factor. J. Anim. Breed. Genet. 135, 263–274. https://doi.org/10.1111/jbg.12335

VanRaden, P.M., 2008. Efficient methods to compute genomic predictions. J. Dairy Sci. 91, 4414–4423. https://doi.org/10.3168/jds.2007-0980

Villanueva, B., Pong-Wong, R., Fernández, J., Toro, M.A., 2005. Benefits from marker-assisted selection under an additive polygenic genetic model1. J. Anim. Sci. 83, 1747–1752. https://doi.org/10.2527/2005.8381747x

Wientjes, Y.C.J., Veerkamp, R.F., Calus, M.P.L., 2013. The Effect of Linkage Disequilibrium and Family Relationships on the Reliability of Genomic Prediction. Genetics 193, 621–631. https://doi.org/10.1534/genetics.112.146290

Yáñez, J.M., Lhorente, J.P., Bassini, L.N., Oyarzún, M., Neira, R., Newman, S., 2014a. Genetic covariation between resistance against both Caligus rogercresseyi and Piscirickettsia salmonis, and body weight in Atlantic salmon (Salmo salar). Aquaculture 433, 295–298. https://doi.org/10.1016/j.aquaculture.2014.06.026

Yáñez, J.M., Naswa, S., López, M.E., Bassini, L., Cabrejos, M.E., Gilbey, J., Bernatchez, L., Norris, A., Soto, C., Eisenhart, J., Simpson, B., Neira, R., Lhorente, J.P., Schnable, P., Newman, S., Mileham, A., Deeb, N., 2014b. Development of a 200K SNP Array for Atlantic Salmon: Exploiting Across Continents Genetic Variation. Proc. World Congr. Genet. Appl. Livest. Prod. Species Breeding: Breeding in aquaculture species, 263.

Yáñez, J.M., Naswa, S., López, M.E., Bassini, L., Correa, K., Gilbey, J., Bernatchez, L., Norris, A., Neira, R., Lhorente, J.P., Schnable, P.S., Newman, S., Mileham, A., Deeb, N., Di Genova, A., Maass, A., 2016. Genomewide single nucleotide polymorphism discovery in Atlantic salmon (Salmo salar): validation in wild and farmed American and European populations. Mol. Ecol. Resour. 16, 1002–1011. https://doi.org/10.1111/1755-0998.12503

Yang, J., Lee, S.H., Goddard, M.E., Visscher, P.M., 2011. GCTA: A Tool for Genome-wide Complex Trait Analysis. Am. J. Hum. Genet. 88, 76–82. https://doi.org/10.1016/j.ajhg.2010.11.011

Yoshida, G.M., Bangera, R., Carvalheiro, R., Correa, K., Figueroa, R., Lhorente, J.P., Yáñez, J.M., 2018a. Genomic Prediction Accuracy for Resistance Against Piscirickettsia salmonis in Farmed Rainbow Trout. G3 Bethesda Md 8, 719–726. https://doi.org/10.1534/g3.117.300499

Yoshida, G.M., Carvalheiro, R., Lhorente, J.P., Correa, K., Figueroa, R., Houston, R.D., Yáñez, J.M., 2018b. Accuracy of genotype imputation and genomic predictions in a two-generation farmed Atlantic salmon population using high-density and low-density SNP panels. Aquaculture 491, 147–154. https://doi.org/10.1016/j.aquaculture.2018.03.004

Yoshida, G.M., Lhorente, J.P., Carvalheiro, R., Yáñez, J.M., 2017. Bayesian genome-wide association analysis for body weight in farmed Atlantic salmon (Salmo salar L.). Anim. Genet. 48, 698–703. https://doi.org/10.1111/age.12621

Yoshida, G.M., Yáñez, J.M., 2021. Increased accuracy of genomic predictions for growth under chronic thermal stress in rainbow trout by prioritizing variants from GWAS using imputed sequence data. Evol. Appl. n/a. https://doi.org/10.1111/eva.13240

You, X., Shan, X., Shi, Q., 2020. Research advances in the genomics and applications for molecular breeding of aquaculture animals. Aquaculture 526, 735357. https://doi.org/10.1016/j.aquaculture.2020.735357

Zenger, K.R., Khatkar, M.S., Jones, D.B., Khalilisamani, N., Jerry, D.R., Raadsma, H.W., 2019. Genomic Selection in Aquaculture: Application, Limitations and Opportunities With Special Reference to Marine Shrimp and Pearl Oysters. Front. Genet. 9. https://doi.org/10.3389/fgene.2018.00693

