## Supplementary Tables for "The impact of genetic relationship between training and validation populations on genomic prediction accuracy in Atlantic salmon"

**Supplementary Material**

**Supplementary Table S1. Number of SNP selected for the corresponding density panels**

| **Density** | **Number of Selected SNPs** |
| --- | --- |
| 100 | 115 |
| 300 | 312 |
| 500 | 512 |
| 800 | 815 |
| 1,000 | 1,012 |
| 3,000 | 3,013 |
| 5,000 | 5,013 |
| 10,000 | 10,005 |
| 20,000 | 19,819 |

**Supplementary Table S2. Bias of GEBVs for decreasing density of SNP panels**

| **SNP Density panel** | **SLC** | **BW** |
| --- | --- | --- |
| 100 | 0.97 ± 0.349 | 1.02 ± 0.294 |
| 300 | 1.01 ± 0.287 | 1.03 ± 0.252 |
| 500 | 0.99 ± 0.266 | 1.02± 0.197 |
| 800 | 1.00 ± 0.250 | 1.03 ± 0.185 |
| 1,000 | 1.02 ± 0.245 | 1.03 ± 0.191 |
| 3,000 | 1.00 ± 0.225 | 1.02 ± 0.172 |
| 5,000 | 0.98 ± 0.214 | 1.02 ± 0.166 |
| 10,000 | 0.98 ± 0.210 | 1.03 ± 0.161 |
| 20,000 | 0.99 ± 0.203 | 1.02 ± 0.160 |

SLC = sea lice count, BW = body weight measured at tagging for 2010 and prior to infection for 2014.

Bias= regression coefficient of (phenotype ~ prediction)

Mean ± standard deviation of bias over 5 folds and 10 cross validation sets.

**Supplementary Table S3. Bias of GEBVs for decreasing genomic relationship between training and validation populations.**

| **Kinship threshold** | **SLC** | **BW** |
| --- | --- | --- |
| 0.3 | 0.73 ± 0.231 | 0.69 ± 0.232 |
| 0.33 | 0.84 ± 0.165 | 0.99 ± 0.170 |
| 0.35 | 0.91 ± 0.304 | 1.02 ± 0.225 |
| 0.37 | 0.92 ± 0.126 | 1.06 ± 0.111 |
| 0.4 | 0.98 ± 0.137 | 1.04 ± 0.099 |
| 0.45 | 0.99 +/ 0.130 | 1.02 ± 0.139 |
| 0.5 | 0.99 ± 0.154 | 1.03 ± 0.119 |
| 0.55 | 0.99 ± 0.135 | 1.04 ± 0.105 |
| No threshold | 1.00 ± 0.146 | 1.04 ± 0.100 |

SLC = sea lice count, BW = body weight measured at tagging for 2010 and prior to infection for 2014.

Bias= regression coefficient of (phenotype ~ prediction)

Mean ± standard deviation of bias over 5 folds and 10 cross validation sets

**Supplementary Table S4A. Accuracy of GEBVs for decreasing SNP density and genomic relationship between training and validation population for sea lice count.**

| SNP density  Kinship  threshold | **100 SNP** | **500 SNP** | **1,000 SNP** | **5,000 SNP** | **10,000 SNP** |
| --- | --- | --- | --- | --- | --- |
| **0.3** | 0.06 ± 0.088 | 0.14 ± 0.111 | 0.20 ± 0.079 | 0.26 ± 0.096 | 0.29 ± 0.094 |
| **0.33** | 0.14 ± 0.081 | 0.28 ± 0.073 | 0.33 ± 0.049 | 0.43 ± 0.059 | 0.45 ± 0.052 |
| **0.35** | 0.17 ± 0.069 | 0.30 ± 0.064 | 0.37 ± 0.072 | 0.46 ± 0.067 | 0.49 ± 0.070 |
| **0.37** | 0.23 ± 0.065 | 0.39 ± 0.061 | 0.46 ± 0.053 | 0.55 ± 0.048 | 0.58 ± 0.049 |
| **0.4** | 0.25 ± 0.061 | 0.41 ± 0.056 | 0.48 ± 0.041 | 0.57 ± 0.038 | 0.59 ± 0.037 |

Mean ± standard deviation of accuracy estimated as correlation(phenotype, GEBV)/h_g_ over 5 folds and 10 cross validation sets

**Supplementary Table S4B. Accuracy of GEBVs for decreasing SNP density and genomic relationship between training and validation population for body weight.**

| SNP density  Kinship  threshold | **100 SNP** | **500 SNP** | **1,000 SNP** | **5,000 SNP** | **10,000 SNP** |
| --- | --- | --- | --- | --- | --- |
| **0.3** | 0.07 ± 0.089 | 0.15 ± 0.117 | 0.20 ± 0.086 | 0.26 ± 0.104 | 0.30 ± 0.103 |
| **0.33** | 0.14 ± 0.080 | 0.27 ± 0.077 | 0.32 ± 0.048 | 0.42 ± 0.053 | 0.45 ± 0.047 |
| **0.35** | 0.17 ± 0.067 | 0.30 ± 0.064 | 0.37 ± 0.070 | 0.46 ± 0.063 | 0.48 ± 0.067 |
| **0.37** | 0.23 ± 0.064 | 0.39 ± 0.062 | 0.45 ± 0.055 | 0.54 ± 0.050 | 0.57 ± 0.052 |
| **0.4** | 0.25 ± 0.060 | 0.40 ± 0.057 | 0.47 ± 0.041 | 0.56 ± 0.038 | 0.58 ± 0.037 |

Mean ± standard deviation of accuracy estimated as correlation(phenotype, GEBV)/h_g_ over 5 folds and 10 cross validation sets

**Supplementary Table S5A. Prediction bias of GEBVs for decreasing SNP density and genomic relationship between training and validation population for sea lice count.**

| SNP density  Kinship  threshold | **100 SNP** | **500 SNP** | **1,000 SNP** | **5,000 SNP** | **10,000 SNP** |
| --- | --- | --- | --- | --- | --- |
| **0.3** | 0.48 ± 0.320 | 0.45 ± 0.209 | 0.58 ± 0.251 | 0.68 ± 0.214 | 0.70 ± 0.220 |
| **0.33** | 0.69 ± 0.296 | 0.72 ± 0.256 | 0.82 ± 0.256 | 0.83 ± 0.188 | 0.84 ± 0.181 |
| **0.35** | 0.75 ± 0.422 | 0.79 ± 0.326 | 0.88 ± 0.317 | 0.90 ± 0.338 | 0.89 ± 0.311 |
| **0.37** | 0.89 ± 0.259 | 0.90 ± 0.189 | 0.93 ± 0.180 | 0.91 ± 0.139 | 0.91 ± 0.133 |
| **0.4** | 0.98 ± 0.234 | 0.97 ± 0.190 | 1.03 ± 0.178 | 0.98 ± 0.142 | 0.98 ± 0.141 |

Mean ± standard deviation of prediction bias over 5 folds and 10 cross validation sets

**Supplementary Table S5B. Prediction bias of GEBVs for decreasing SNP density and genomic relationship between training and validation population for body weight.**

| SNP density  Kinship  threshold | **100 SNP** | **500 SNP** | **1,000 SNP** | **5,000 SNP** | **10,000 SNP** |
| --- | --- | --- | --- | --- | --- |
| **0.3** | 0.25 ± 0.336 | 0.39 ± 0.324 | 0.52 ± 0.217 | 0.63 ± 0.249 | 0.70 ± 0.235 |
| **0.33** | 0.55 ± 0.286 | 0.74 ± 0.187 | 0.80 ± 0.143 | 0.93 ± 0.179 | 0.97 ± 0.161 |
| **0.35** | 0.71 ± 0.285 | 0.99 ± 0.183 | 0.90 ± 0.229 | 0.98 ± 0.208 | 1.01 ± 0.217 |
| **0.37** | 0.91 ± 0.215 | 1.01 ± 0.142 | 1.03 ± 0.144 | 1.05 ± 0.121 | 1.07 ± 0.116 |
| **0.4** | 0.99 ± 0.225 | 1.02 ± 0.152 | 1.05 ± 0.126 | 1.04 ± 0.108 | 1.05 ± 0.102 |

Mean ± standard deviation of prediction bias over 5 folds and 10 cross validation sets

**Supplementary Figure S1. Accuracy for different density of SNPs panels**

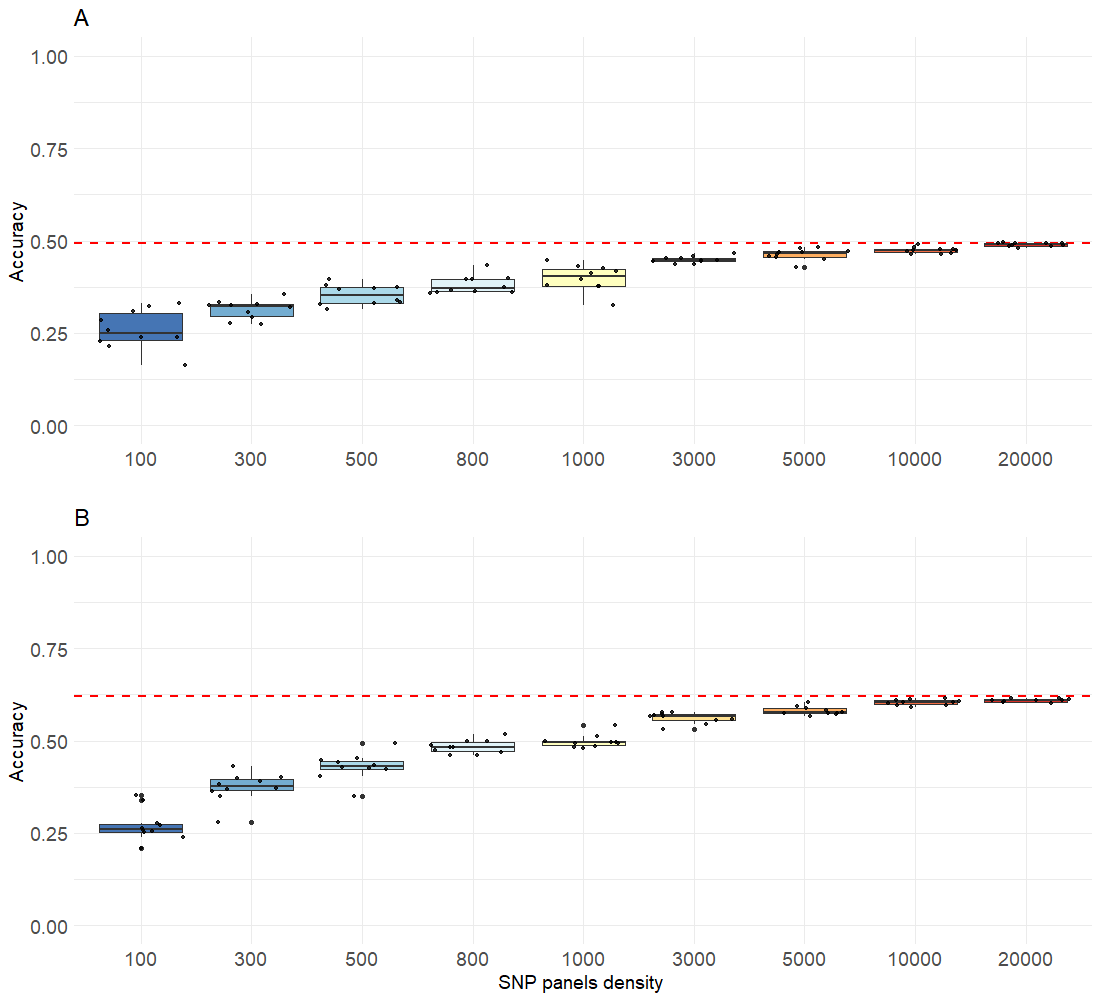

Box plot of accuracy of genomic prediction (GBLUP) for sea lice count (A) or initial body weight (B) estimated using various density SNPs panels. For each density panel, 10 replicates were used and genomic prediction were averaged over 5 folds and 10 cross validation sets. The red dotted line correspond to the mean of genomic prediction obtained with the fully imputed dataset (32K SNPs).
